# Birth weight associations with psychiatric and physical health, cognitive function, and DNA methylation differences in an adult population

**DOI:** 10.1101/664045

**Authors:** Rebecca A. Madden, Daniel L. McCartney, Rosie M. Walker, Robert F. Hillary, Mairead L. Bermingham, Konrad Rawlik, Stewart W. Morris, Archie Campbell, David J. Porteous, Ian J. Deary, Kathryn L. Evans, Jonathan Hafferty, Andrew M. McIntosh, Riccardo E. Marioni

## Abstract

**Background:** The Developmental Origins of Adult Disease (DOAD) theory predicts that prenatal and early life events shape adult health outcomes. Birth weight is a useful indicator of the foetal experience, and has been associated with multiple adult health outcomes. DNA methylation (DNAm) is one plausible mechanism behind the relationship of birth weight to adult health.

**Methods:** The Generation Scotland study allows data linkage to historic Scottish birth cohorts, and birth records held through the NHS Information and Statistics Division. Data linkage with these sources yielded a sample of 4, 710 individuals. Health measures were related to birth weight in regression models. An epigenome-wide association study (EWAS) was performed in a subgroup (n=1, 395), relating adult DNAm from whole blood to birth weight, with replication in an independent sample (n=362). Associations between birth weight and epigenetic clocks were also assessed.

**Findings:** Higher birth weight was significantly associated with reduced incidence of depression and osteoarthritis, higher body mass index, and higher general intelligence (absolute standardised effect size range 0·04 to 0·30, p_(FDR)_<0·05). Meta-analysis of discovery and replication EWAS studies yielded one genome-wide significant CpG site (p=5·97×10^−9^), cg00966482. Significant associations between birth weight and Grim Age (p=0·0014) and DNAm-derived telomere length (p=3·3×10^−4^) are also described.

**Interpretation:** Our results demonstrate associations between birth weight and adult health outcomes, with particularly striking effects for depression risk. It also provides support for an association between birth weight and DNAm, describing the first significant EWAS site associated with birth weight in an adult sample.

**Funding:** Wellcome Trust Strategic Award 104036/Z/14/Z

**Research in Context:** *Evidence before this study:* The associations between birth weight and various adult health outcomes have been well established. DNA methylation is a plausible mechanism through which early life experiences may continue to affect health throughout the lifecourse; however, evidence for birth weight associations with DNA methylation in adulthood has not yet been robustly established. This is likely due to small sample sizes of previous samples, as well as the use of poor-quality birth weight data, such as binary ‘low/normal’ variables or retrospective self-report. Alternatively, work has attempted to describe the persistence into adulthood of DNA methylation at sites identified at birth.

*Added value of this study:* We investigated genome-wide differential DNA methylation patterns from whole blood using data linkage-derived, continuous birth weight data, in the largest reported adult sample (n=1, 395) with replication (n=362) and meta-analysis. Meta-analysis revealed one epigenome-wide significant CpG site, to our knowledge the first significant EWAS result reported for birth weight in a an adult sample. In addition, we found associations between birth weight and GrimAge and a DNA methylation-derived measure of telomere length, demonstrating accelerated biological ageing in lower birth weight individuals. Together, these results suggest differential methylation exists in adulthood related to birth weight, and this may be relevant to health and mortality.

*Implications of all the available evidence:* Although CpG sites differentially methylated with birth weight at parturition may not remain so throughout life, the adult epigenome may still provide information on the impact of birth weight on health outcomes. The adult epigenome, therefore, may represent a useful archive of the foetal experience which results in birth weight variability, and this information may provide clinically useful information in mid-life.

## Introduction

The Developmental Origins of Adult Disease theory (DOAD) states that through developmental plasticity, the foetal experience can permanently influence adult health (1). The theory’s main proponent, David Barker, originally relied on birth weight as an index of foetal nutrition – an assumption that has been contested by the awareness that multiple factors can influence birth weight (2). Maternal stress, illness, and socioeconomic status (3-5) are among modifiable influences over offspring birth weight; in addition, maternal-specific genetic variation has been found to influence birth weight, acting through the intrauterine environment (6). Thus, birth weight can be seen as an index of the general foetal experience.

There is evidence of a U-shaped association between birth weight and adult health. Birth weight is clinically conceptualised as ‘low’ below 2·5kg, and ‘high’ above 4·5kg (7), however it can also be analysed on a continuous scale. Lower birth weight has been associated with an increased risk of heart disease, type II diabetes, stroke, and hypertension (8-11). Low birth weight is also associated with poorer cognitive ability, and a raised risk for mood disorders (12, 13). At the higher end of the spectrum, birth weight is associated with higher body mass index, and higher risk of breast cancer (14, 15). These associations are found after accounting for adult lifestyle factors, such as smoking and BMI, indicating a residual effect of birth weight on health outcomes. Since birth weight information is recorded as standard practice, it could be considered a useful and readily available predictive tool for adult health risks.

The DOAD theory is a plausible explanation for these observations. Prenatal factors affect the foetus in its highly plastic state, giving rise to birth weight variability, and also to developmental changes which permanently affect the function and health of organs and systems (1). Some of these changes may be structural, affecting for example, vascular development and function (16). DNA methylation (DNAm) is an epigenetic modification that can be influenced by genetics or by environmental factors throughout life. CpG methylation is characterised by the addition of a methyl group to cytosine nucleotides in the context of cytosine-guanine dinucleotides. DNAm changes are linked to the regulation of gene expression, providing a possible mechanism through which environmental influences may have lasting biological effects (17). Therefore, DNAm is one putative mechanism through which developmental experience may influence adult health.

Birth weight effects on the epigenome have been previously described in cord blood (18, 19) and during childhood (20), seeming to diminish into adolescence and beyond (18, 21). It is possible, however, that examining only the persistence of DNAm patterns associated with birthweight at the point of birth ignores the full adult epigenome which, despite having changed since birth, nevertheless may provide information on the impact of birthweight on downstream health. Here, we perform the first large EWAS study of birth weight on adult DNAm. In addition, recent work has demonstrated the utility of so-called “epigenetic clocks” in demonstrating elements of vulnerability to disease and mortality (22), conceptualised as accelerated biological age in comparison to chronological age. These analyses together have the potential to describe the known associations between birthweight and many aspects of whole-body health in adulthood.

Here, we describe associations between birth weight and a range of adult health conditions and risk factors, using high-quality data that improves on previous work in the field (e.g. (13)); it also reports an Epigenome-Wide Association Study (EWAS) of birthweight using whole-blood DNA derived in an adult sample, and associations between birth weight and five epigenetic clocks. While previous work has looked at DNAm relationships to birth weight in adults, none have performed EWAS studies in this age group, looking instead at persistence of DNAm effects from birth (18, 21). We, therefore, hypothesise that while birth weight associated DNAm patterns may change over time, differences will still exist in adulthood.

## Materials and Methods

### Generation Scotland and other Cohorts

Generation Scotland (GS) is a Scottish family-based cohort n=23, 690 (23). Data were collected from participants between 2006 and 2011. GS is a deeply-phenotyped cohort, allowing examination of many aspects of adult health, alongside genomic and biometric measures. In addition, 98% of GS participants gave informed consent for data linkage to routinely collected health data and to information from other Scottish population cohort studies, both current and historical. These include several with neonatal and maternity information: the Aberdeen Children of the 1950s ((24), ACONF); the Aberdeen Maternity and Neonatal Databank ((25), AMND); the Walker Birth Cohort (26); and the Scottish Morbidity Records ((27), SMR02 – the Maternity Inpatient and Day Case record, and SMR11 – the Neonatal Inpatient dataset). Birth weight in grams, alongside gestational age at birth and twin information, was collated from these sources and linked to adult GS records for 4, 713 participants (**Supplementary File 1, Figure 1**). Birth weight data derived from health records taken at birth improves on the self-reported nature of birth weight variables found in other population cohorts.

**Figure 1:**
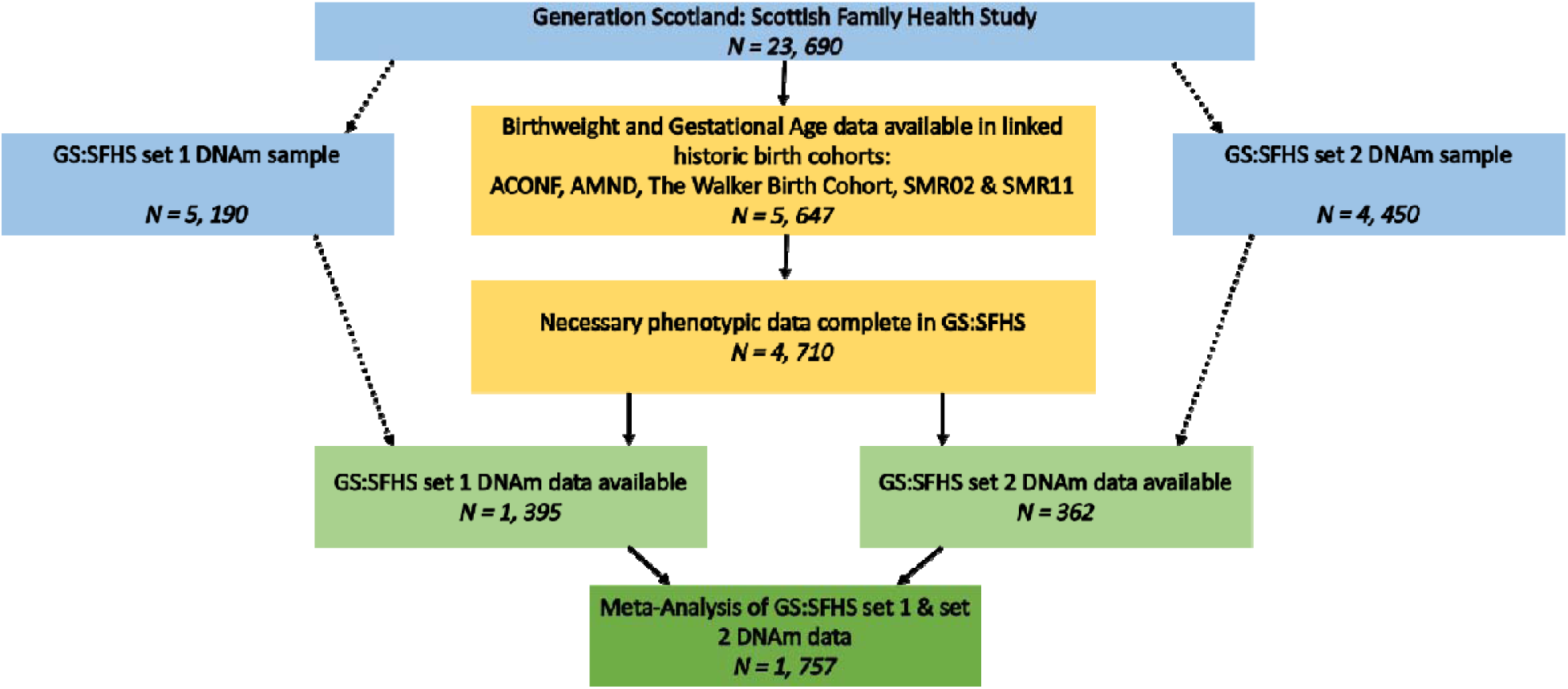
Inclusion flow diagram detailing the selection of samples and subsamples for the current study.

### Statistical Analyses

All analyses were conducted in R version 3·5·1 (28).

To control for the known effects of gestational age and sex on birthweight (29), we considered the residuals from a regression model in place of raw birth weight throughout:

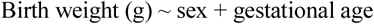

Self-reported diseases (yes/no) that were commonly reported in the cohort (prevalence >1%) included depression (prevalence 6·6%), osteoarthritis (1·6%), high blood pressure (3·4%), diabetes (1·3%), and asthma (16·6%). These were modelled against the birth weight residuals using logistic regression. Average diastolic and systolic blood pressure measurements, body mass index, high-density lipoprotein (HDL) cholesterol, and general intelligence score (derived from multiple cognitive tests - see **Supplementary File 1**) were examined as continuous variables in linear regression models. Depression diagnosed using the Structured Clinical Interview for DSM-IV (SCID;(30) was also examined, as a binary yes/no for a diagnosis of major depressive disorder (**Supplementary File 1**). Three individuals who had responded ‘yes’ to all self-reported illnesses were excluded from analyses, leaving a final population of n=4, 710. Some traits had missing values, which resulted in case-wise exclusion from the relevant regression model (**Table 1**).

**Table 1:** Population characteristics of the phenotyped and EWAS samples.

|  | Phenotype Sample |  |  | EWAS Sample |  |  |
| --- | --- | --- | --- | --- | --- | --- |
|  | N | Mean | SD | N | Mean | SD |
| Age (years) | 4, 710 | 29.5 | 10.9 | 1, 395 | 37.1 | 14.7 |
| Birthweight (g) | 4, 710 | 3, 399 | 516 | 1, 395 | 3, 377 | 518 |
| Gestation (weeks) | 4, 710 | 39.8 | 1.7 | 1, 395 | 40 | 1.8 |
| BMI (kg/m<sup>2</sup>) | 4, 385 | 25.2 | 5.2 | 1, 387 | 26.1 | 5.4 |
| Education* | 4, 411 | 5 | 4-6 | 1, 317 | 5 | 3-6 |
|  | N | % |  | N | % |  |
| Sex - Male | 2, 025 | 43.0 |  | 569 | 40.8 |  |
| Female | 2, 688 | 57.0 |  | 826 | 59.2 |  |
| Socioeconomic Status** | (4, 389) |  |  | (1, 310) |  |  |
| Quintile 1 (most deprived) | 621 | 14.2 |  | 207 | 15.8 |  |
| Quintile 2 | 717 | 16.3 |  | 210 | 16.0 |  |
| Quintile 3 | 715 | 16.3 |  | 183 | 14.0 |  |
| Quintile 4 | 1, 064 | 24.2 |  | 301 | 23.0 |  |
| Quintile 5 (least deprived) | 1, 272 | 29.0 |  | 409 | 31.2 |  |
| Smoking | (4, 525) |  |  | (1, 344) |  |  |
| Current Smoker | 918 | 20.3 |  | 257 | 19.1 |  |
| Ex-Smoker (<12 months) | 237 | 5.2 |  | 56 | 4.2 |  |
| Ex-Smoker (>12 months) | 662 | 14.6 |  | 273 | 20.3 |  |
| Never Smoker | 2, 708 | 59.8 |  | 758 | 56.4 |  |

\* Median and Interquartile range reported. Education was coded as an ordinal variable: 0 = 0yrs, 1 = 1-4yrs, 2 = 5-9yrs, 3 = 10-11yrs, 4 = 12-13yrs, 5 = 14-15yrs, 6 = 16-17yrs, 7 = 18-19yrs, 8 = 20-21yrs, 9 = 22-23yrs, 10 = ≥24yrs.
\*\*SIMD Quintile. SIMD is the Scottish Index of Multiple Deprivation, a postcode-derived index of socioeconomic status. The quintiles derived on the full Generation Scotland cohort ranged from 1 (most deprived) to 5 (least deprived).

Adult health and functional domains were regressed against birth weight residuals and relevant covariates in a simple model:

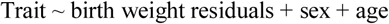

And in a fully-adjusted model:

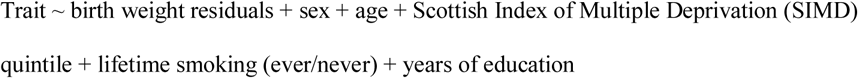

For the general intelligence trait, years of education was removed from this fully-adjusted model as the two are highly collinear (Pearson r=0·36).

Correction for multiple testing was carried out using a false discovery rate P<0·05.

### Epigenome-wide association study

Genome-wide DNA methylation was profiled in 5, 190 individuals in GS, taken from peripheral whole blood, using the Illumina HumanMethylationEPIC BeadChip (Illumina Inc., San Diego, CA). Quality control and normalisation were carried out as described elsewhere ((31, 32); **Supplementary File 2**). For the subgroup of GS for whom birth weight and gestational age information were available, DNAm data was available for n=1, 395 (**Figure 1)**.

In a second release of data generated during this analysis, using a near identical protocol (**Supplementary File 2**), an independent set of methylation data became available for an additional 4, 450 GS participants. In this replication sample, a further 362 participants with both birth weight and gestational age information were used.

The birth weight residuals described above were used in the EWAS model, which was run using the ‘limma’ package in R (empirical Bayes moderated t-statistics). The discovery EWAS model used CpGs corrected for relatedness (**Supplementary File 2**), as the first batch of DNAm data was collected on related individuals:

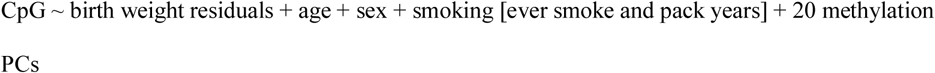

Additional covariates (estimated white blood cell proportions – CD4T, CD8T, Granulocytes, BCells, Natural Killer cells – and methylation batch), which were regressed out during the relatedness pre-correction for the discovery dataset, were also included in the replication dataset of unrelated individuals.

### Meta-Analysis of discovery and replication samples

A standard error-weighted meta-analysis of the discovery and replication EWASs was performed using the METAL software package. Summary statistics from the discovery, replication and meta-analysis EWASs are available at [**weblink to be inserted upon acceptance**].

### Sensitivity analysis

As a sensitivity analysis, the 20 CpGs most strongly associated with birth weight in the discovery EWAS were re-examined with a maximally-corrected model to account for additional lifestyle factors which may affect both DNAm and health. These were body mass index, socioeconomic status (Scottish Index of Multiple Deprivation - SIMD), and number of years of education, on top of the correction for smoking already included in the discovery EWAS model. DNAm data were pre-corrected for batch, cell counts, and relatedness. The maximally corrected model was:

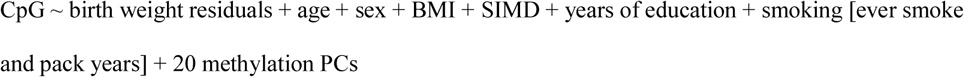

### Epigenetic clock analyses

Detail on the estimation of the five epigenetic clocks used has previously been reported elsewhere (22). Briefly, penalised regression models have used to identify subsets of CpG sites that predict chronological age (33, 34), biological age (35), telomere length (36), and survival (37). The clocks either trained predictors directly on the outcome (e.g., chronological age or telomere length), or via surrogate markers (e.g., protein levels and biomarkers that are known to associated with biological/health processes). The residuals from regressions of these predictors on chronological age gives an index of age acceleration. Here, we considered the association (linear regression) between birth weight (outcome) and the five sets of age acceleration residuals, with covariate adjustment for age, sex, methylation set, and estimated white cell proportions.

### Ethics approval and consent to participate

All components of GS received ethical approval from the NHS Tayside Committee on Medical Research Ethics (REC Reference Number: 05/S1401/89). GS has also been granted Research Tissue Bank status by the Tayside Committee on Medical Research Ethics (REC Reference Number: 10/S1402/20), providing generic ethical approval for a wide range of uses within medical research.

## Results

### Phenotype population characteristics

There were 4,710 Generation Scotland (GS) participants with birth weight and gestational age information available (**Table 1**). The population was 57% female, with a mean birth weight of 3·40kg (SD=0·52), and gestational age of 39·8 weeks (SD=1·7). The minimum birth weight in the sample was 0·7kg, the maximum was 5·1kg. Using the clinical cut-off of 2·5kg, 3·99% (n=188) of the sample were of clinically ‘low’ birth weight. This means the data describe phenotypic relationships to birth weight across pathological and non-pathological cases.

### Relationship of adult outcomes to birth weight

Regression models identified significant associations between higher birth weight (effect sizes are reported per SD) and lower risk of SCID-diagnosed depression (OR=0·86; 95% CI 0·78-0·95, p=0·018); lower risk of self-reported osteoarthritis (OR=0·74; 95% CI 0·58-0·94; p=0·034); higher BMI (β=0·072; SE=0·016, p=3·7×10^−05^) and higher general intelligence (β=0·042; SE=0·016, p=0·027; **Table 2**).

**Table 2:** Outputs of logistic and linear regression models of health traits ∼ birth weight residuals + age + sex + socioeconomic status (SIMD) + ever smoke + education years, with FDR-correction for multiple testing.

| Trait | Odds Ratio | 95% CI | P value | P (FDR) |
| --- | --- | --- | --- | --- |
| SCID depression | 0.86 | 0.78 – 0.95 | 0.0033 | 0.018 |
| SR depression | 0.92 | 0.81 – 1.05 | 0.21 | 0.38 |
| SR hypertension | 0.93 | 0.78 – 1.09 | 0.36 | 0.44 |
| SR diabetes | 0.96 | 0.74 – 1.25 | 0.76 | 0.76 |
| SR osteoarthritis | 0.74 | 0.58 – 0.94 | 0.012 | 0.034 |
| SR asthma | 0.96 | 0.88 – 1.04 | 0.3 | 0.43 |
| Trait | Beta | SE | P value | P (FDR) |
| Body Mass Index (kg/m<sup>2</sup>) | 0.072 | 0.016 | 3.4x10<sup>-6</sup> | 3.7x10<sup>-5</sup> |
| HDL cholesterol | 0.025 | 0.016 | 0.12 | 0.27 |
| Average systolic BP | -0.014 | 0.014 | 0.32 | 0.43 |
| Average diastolic BP | -0.011 | 0.015 | 0.47 | 0.51 |
| General intelligence* | 0.042 | 0.016 | 0.0074 | 0.027 |

\*For the ‘g’ general intelligence factor, education years was removed from the model as it resulted in over-correction of the effect.

In minimally-adjusted regression models (covarying for age and sex alone), higher birth weight showed a relationship with lower self-reported depression risk (OR=0·85; 95% CI 0·75-0·96; p=0·018). Associations were also found for the four traits described above, but no other adult health or functional outcomes (**Supplementary Table 1**).

### Epigenome-wide association study of birthweight

The epigenome-wide association study of birth weight revealed no CpGs significant at the genome-wide level (p<3·6×10^−8^; (38)), although 19 CpGs had P<1×10^−5^ (minimum FDR corrected P-value of 6·05×10^−8^ for cg00966482) (**Figure 2a; Supplementary Table 2**). The 19 CpGs were largely uncorrelated, with the exception of three CpGs (located within *CASZ1* – two within 200 base pairs of each other, with the third site around 11kb away – **Supplementary Table 2**) that had absolute r≥0·6(**Figure 3**).

**Figure 2:**
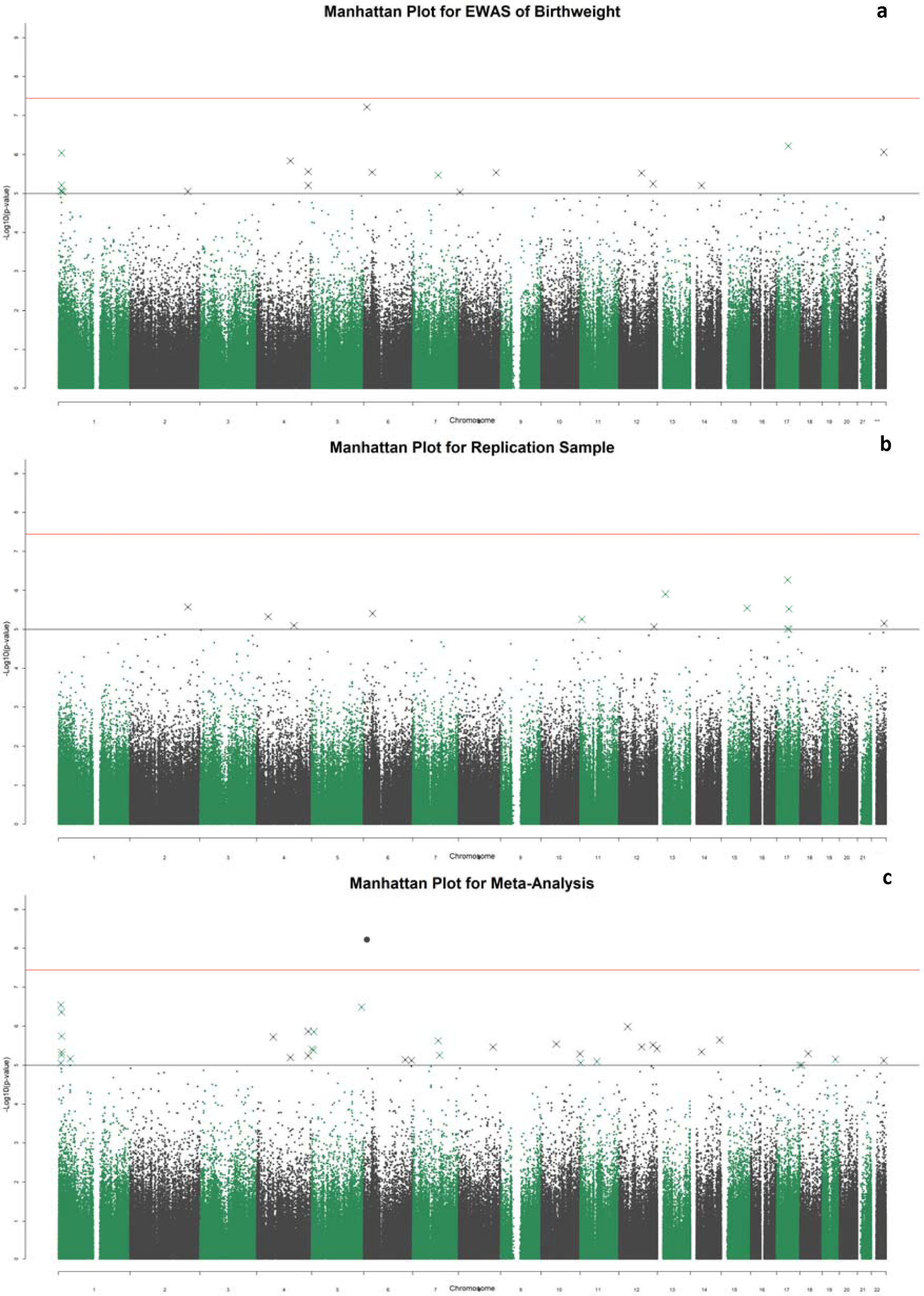
Manhattan plots for the epigenome-wide association study of birth weight (**a**); the replication sample EWAS (**b**); and the meta-analysis of the main EWAS and the replication (**c**). The black and red lines represent the suggestive, and Bonferroni corrected P-value thresholds of P=1×10^−5^ and 3·6×10^−8^, respectively.

**Figure 3:**
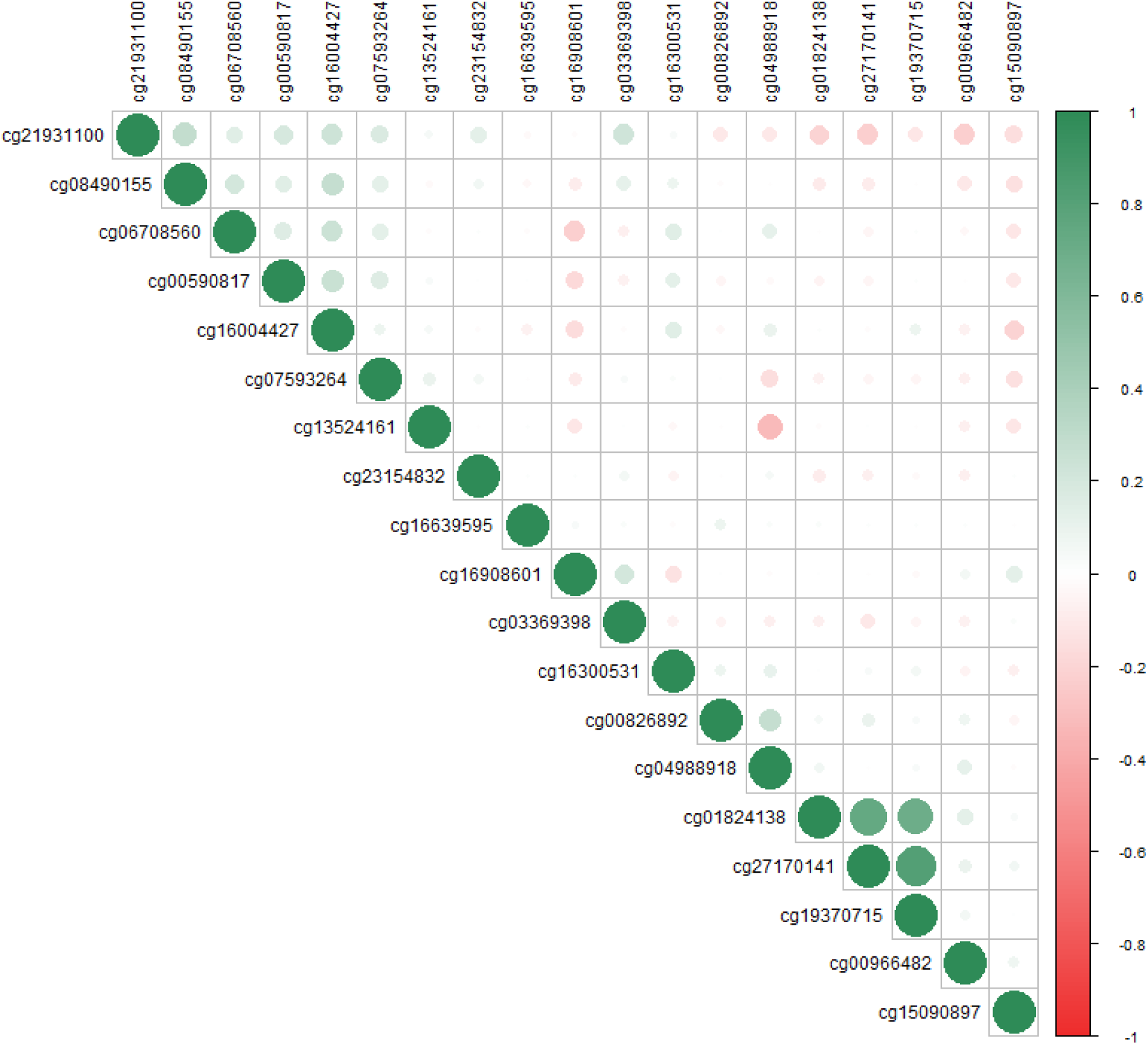
Correlation plot between methylation at the top 19 CpG sites from the epigenome-wide association study. The shade and scale of the dots represent the magnitude and direction of the correlation between pairs of CpGs.

### Replication in an independent DNAm sample

There were no genome-wide significant associations in the replication sample (n=362) (**Figure 2b; Supplementary Table 3**).

Of the 19 CpGs that exceeded the suggestive significance threshold in the discovery EWAS, two reached nominal significance (p<0·05) in the replication analysis: cg00590817 (p=0·0069), and cg00966482 (p=0·031). The latter CpG was the most significantly associated site from the discovery EWAS (p=6·05×10^−8^) and is found within the *HERV-FRD* (also referred to as *ERVFRD-1*) gene. There was good concordance between the effect sizes of DNAm associations with birthweight in the discovery and replication studies (r=0·59) (**Figure 4**).

**Figure 4:**
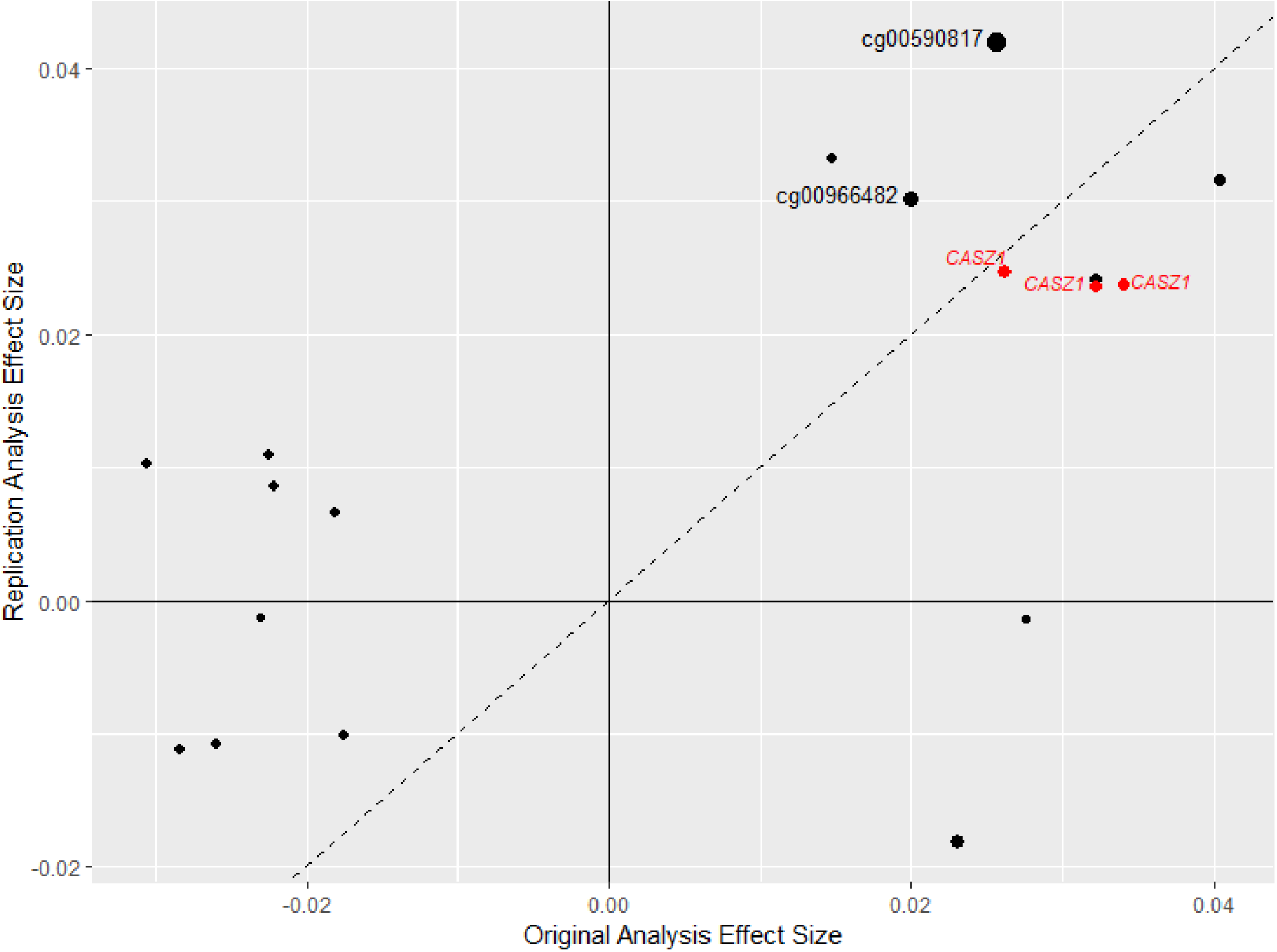
Effect sizes for 18 of the top 19 CpG sites in the discovery sample plotted against the effect sizes in the replication sample (cg04988918 was not included in the replication array). The point size is determined by the −log10 of the p-values for these hits in the replication analysis. The two points labelled in black are the two CpG sites which achieved nominal significance in the replication study, and the three highlighted in red are the three co-methylated CpG sites within the *CASZ1* gene.

### Meta-analysis of discovery and replication samples

Meta-analysis of the original discovery EWAS sample with the replication sample found a genome-wide significant effect of birthweight on DNAm at the CpG site cg00966482 located in the *ERVFRD-1* gene (p=5·97×10^−9^; **Figure 2c**; **Supplementary Table 4**).

### Adjusting the EWAS model for lifestyle factors

As a sensitivity analysis, we re-ran the EWAS pipeline for the top CpG sites, with additional corrections for BMI, education, and SIMD. This resulted in some minor changes to the effect sizes (mean attenuation of 1·35%, range 11·12% attenuation to 12·67% increase in effect size; **Supplementary Table 5**).

### Relationship of birth weight to epigenetic signatures of age and telomere length

The mean values for the five epigenetic clocks were: Horvath age 44·7 yrs (SD=10·6); Hannum age 36·1 yrs (SD=10·2); PhenoAge 32·8 yrs (SD=12); GrimAge 45·7 yrs (SD=14·1); DNAm telomere length 7·6 kilobase pairs (SD=0·29). Linear regression models of birth weight residuals (predictor) against five signatures of epigenetic age acceleration revealed significant associations between higher birth weight and both lower Grim Age (β=-0·083; SE=0·026, p=0·0014) and shorter DNAm telomere length (β=0·098; SE=0·027, p=3·3×10^−4^; **Table 3**). Associations between birth weight and the other three measure of epigenetic age acceleration were non-significant.

**Table 3:** Outputs of linear regression models between birth weight residuals and five epigenetic signatures of accelerated biological ageing – Horvath, Hannum, PhenoAge, GrimAge, and DNAmTL.

| Clock | Beta | SE | P value |
| --- | --- | --- | --- |
| Horvath | 0.0032 | 0.025 | 0.9 |
| Hannum | -0.0062 | 0.027 | 0.82 |
| PhenoAge | -0.042 | 0.027 | 0.12 |
| GrimAge | -0.083 | 0.026 | 0.0014 |
| DNAmTL | 0.098 | 0.027 | $3.3 \times 10^{-4}$ |

## Discussion

We observed a strong association between birth weight and adult depression, where a 1 standard deviation (SD; 0·52kg) increase in birth weight was linked to 15% lower odds of future depression. A 1SD increase in birth weight was also linked to a 0·072 SD (∼0·37kg/m^2^) higher adult BMI, a 0·042 SD higher general intelligence score, and 30·7% lower odds for self-reported osteoarthritis. We also identified one genome-wide significant association between birth weight and blood-based DNA methylation in adulthood.

Previous studies have found both positive and negative associations between birth weight and depression. A meta-analysis of 18 studies (n=∼50, 000) found support for a modest effect of low birthweight (<2·5kg) on depression risk (OR=1·15, 95% CI=1·00-1·32), but suggested this was a result of publication bias (13). We improved upon the design of many of the papers within that meta-analysis by utilising clinician-diagnosed depression and birth weight as a continuous variable (compared to self-reported measures of depression or psychological distress and use of a binary (<2·5kg vs normal) birth weight variable). Compared to SCID-diagnosed depression, we found self-reported depression had a weaker association with birth weight. Furthermore, the association with the self-reported measure was statistically significant in the minimally-adjusted but not fully-adjusted regression models. This emphasises the importance of using high quality, clinically meaningful tools for measuring psychiatric disorders. The aetiology of depression is complex, with multiple biological and psychosocial factors contributing increments of risk, thought to sum towards illness. This study indicates that birth weight may be considered as one of these contributing risk factors, as its association with adult diagnosis remained after accounting for many adult lifestyle factors.

Consistent with the published literature (11, 14), we identified an association between higher birth weight and higher adult BMI. Previous work has used Mendelian randomisation techniques to show that birthweight is a causal factor in determining adult BMI (9). We also found a significant relationship between higher birth weight and lower prevalence of osteoarthritis in this sample. Developmental origins of osteoarthritis have been described before, with higher disease prevalence in low birth weight individuals (39), and those with low weight at age one (40).

A positive association between birth weight and general intelligence is also reported here (1SD increase in birthweight results in a 0·042 SD higher general intelligence score). Impaired cognitive ability after low birth weight has been shown in several samples in childhood (12) and into adulthood (41).

Some previously-established effects of birth weight on adult health are not supported here, for instance, links to type 2 diabetes and cardiovascular disease (9). This may be partly due to the self-report nature of the health measures used. Our sample size was also smaller than previous investigations into birth weight and cardiovascular outcomes (9, 11).

In the EWAS meta-analysis we observed a genome-wide significant association between higher birthweight and higher methylation levels at cg00966482 (*ERVFRD-1*). *ERVFRD-1* encodes syncitin-2, a protein involved in placental embedding (42). This site was not identified in a previous meta-analysis EWAS of birthweight (21). Moreover, this is the first report of any genome-wide significant EWAS association for birth weight in an adult sample. DNAm data was derived from whole blood taken during adulthood, so our findings might not generalise to other tissue types. However, there are many advantages to interrogating blood-based methylation levels: first, it is the only tissue type that is readily available in large epidemiological studies – it would be expensive and invasive to obtain adipose tissue samples on over 1, 000 individuals; second, it is systemic and tracks multiple biological processes including biomarkers of inflammation, cardiovascular disease, cardiometabolic disease, all of which are relevant processes related to birthweight; third, the DOAD theory posits that birth weight is related to health outcomes across the whole body.

Of the 19 CpG sites with P<1×10^−5^ in the discovery EWAS, 10 were located within known genes. Some of these genes contain SNPs that have genome-wide associations (with P<5×10^−8^) with cardiovascular, psychiatric, and developmental pathways (**Supplementary Table 6**). Three of the 19 nominally significant CpG sites identified in the discovery EWAS were highly correlated (min r=0·69). Higher birthweight was associated with higher methylation levels at these sites, which were located within *CASZ1*, a gene encoding the zinc finger protein castor homolog 1, a transcriptional activator involved in vascular morphogenesis (43). A differentially methylated region (DMR) in the *CASZ1* gene was recently identified in placental tissue between infants born small vs. large for gestational age (19), thus this study suggests the persistence of differential methylation of this gene into adulthood. Genetic variants in *CASZ1* have previously been implicated in GWAS studies on various aspects of cardiovascular health (**Supplementary Table 6**). These have included studies in multi-ethnic populations on blood pressure (44, 45), and on other cardiovascular health issues such as atrial fibrillation (46) and stroke (47).

A recent longitudinal meta-analysis (21) of DNAm relationships to birth weight identified 914 CpG sites associated with birth weight in 24 EWAS studies from neonatal blood (total n=8, 825). The persistence of methylation differences at these sites was then examined in other cohort data from childhood (total n=2, 756 from 10 studies; 2-12y), adolescence (total n=2, 906 from 6 studies; 16-18y), and adulthood (1, 616 from 3 studies; 30-45y). Nominally significant methylation differences were found to persist into childhood, adolescence, and adulthood at 87, 49, and 42 sites respectively. This supports evidence that birth weight-related methylation differences may attenuate over time (18, 20). It has, however, been demonstrated that some methylation patterns persist into late adulthood after prenatal famine exposure (48). It is therefore plausible that other prenatal factors may continue to affect DNAm into adulthood. This study adds to the available literature, demonstrating that DNAm differences may exist in adulthood which are not persistent from birth, but which nonetheless have the potential to describe part of the ongoing association between birth weight and adult health. Whereas there was good concordance in effect sizes for the top discovery EWAS findings in our replication dataset, none were significant after correction for multiple testing. Further EWAS studies of birth weight and DNAm in mid-life are therefore required to replicate our genome-wide significant finding from the combined discovery and replication meta-analysis.

Significant associations are demonstrated between birth weight and epigenetic predictors of ageing, mortality, and cellular senescence. Higher birth weight is associated with lower GrimAge acceleration – a measure of epigenetic age with validated relationships to health and all-cause mortality (22) – and longer telomeres as estimated by DNAm data, known to associate with a variety of health outcomes (36). These findings further validate the rationale behind the current study, by indicating longitudinal associations between birth weight and DNAm predictors of health in mid-life.

This study exploited rarely available data linkage capacity to acquire neonatal information from birth medical records. The phenotyping and DNAm data available within the cohort have allowed the association of both health traits and DNAm data with birth weight. There are, however, some limitations inherent to the cross-sectional design of this study. Longitudinal data would allow analysis of the persistence of DNAm signatures across time. In addition, the number of individuals for whom accurate birth weight and gestational information could be identified in historical health cohorts limited the sample size for the EWAS analyses.

This study presents the first epigenome wide association study of birth weight on DNA methylation in adulthood, alongside associations between birthweight and both epigenetic age acceleration and telomere length. We also present a comprehensive study of birth weight in relation to cognitive, psychiatric, and disease traits in mid-life. The Developmental Origins of Adult Disease theory predicts that birth weight can affect many domains of adult physical and psychological health. We found evidence to support this at both the molecular and broad disease/health phenotype levels.

## Abbreviations

DOAD: Developmental Origins of Adult Disease
DNAm: DNA methylation
EWAS: epigenome-wide association study
BMI: body mass index
GS: Generation Scotland
OR: Odds Ratio
SD: Standard Deviation
SE: Standard Error
CI: confidence interval
SCID: Structured Clinical Interview for DSM-IV
ACONF: Aberdeen Children of the 1950s
AMND: Aberdeen Maternity and Neonatal Databank
SMR: Scottish Morbidity Records
HDL: high-density lipoprotein

## Acknowledgements

The authors thank all individuals and project team members who have contributed to both GS:SFHS and to the ‘STRADL: Stratifying Resilience and Depression Longitudinally’ follow-up study.

## Author contributions

Conception and design: RAM, JH, AMM, and REM. Data analysis: RAM and DLM. Interpretation: RAM, DLM, JH, AMM, REM. Drafting the article: RAM. Revision of the article: all authors.

## Availability of data and materials

According to the terms of consent for Generation Scotland participants, access to data must be reviewed by the Generation Scotland Access Committee. Applications should be made to. Summary statistics for the EWAS models will be made available on the University of Edinburgh DataShare facility upon acceptance **[doi to be added]**

