## Supplementary material for "Birth weight associations with psychiatric and physical health, cognitive function, and DNA methylation differences in an adult population": Supplemetary Files

**Supplementary File 1:** Identification and calculation of variables in GS.

Identifying Birthweight Information:

Birthweight in Grams, alongside Gestational Age and twin information was merged into GS, using a linker file containing the respective identifiers for each cohort, and any matching GS IDs.

In the Walker record, some birthweight information was stored only in pounds and ounces. For these data points, the following calculation was used to impute birthweight in grams:

(no.lbs/453·59) + (no.oz/28·35) = birthweight in grams

In SMR02, data was stored under the identifier of the mother, rather than the child. The child’s sex, date of birth, and the mother’s GS ID were used to create a ‘key’ in the format “Maternal ID*sex*DOB”, making a unique identifier for each individual allowing for the record to be merged.

Different-sex twin pairs in SMR02 were separated by birth order, and merged with GS in the same manner as singleton births. Same sex twin pairs were excluded if the birthweight discordance exceeded 1SD of the group birthweights. For those remaining, one of each twin pair was randomly excluded – therefore for all same sex twins, one twin’s birthweight info was linked to one adult GS record. In SMR 11, a similar process was undertaken, as some same-sex twin pairs were recorded under identical ID numbers. Again, one twin from each pair was randomly excluded. 19 twin pairs in SMR11, and 24 in SMR02 had their data imputed in this way.

Calculation of General Intelligence Scores:

Cognitive testing was performed by nurses who had been trained and validated by a psychologist. Cognitive function measurements taken in GS [1] included:

-	Logical Memory (sum of immediate and delayed recall) from the Wechsler Memory Scale III

-	The Digit Symbol substitution test from the Wechsler Adult Intelligence Scale III

-	The phonemic verbal fluency test, in which participants listed words beginning with C, F, and L for one minute each.

-	Mill Hill Vocabulary Scale

A general intelligence score was calculated using principal components analysis (PCA). Score from the first unrotated component were extracted from a PCA of the participants’ responses on the Weschler Logical Memory items, the Weschler Digit Symbol task, and the Verbal Fluency test, and the Mill Hill Vocabulary test. Scores greater or less than 3.5 standard deviations from the mean on any test were excluded from this analysis.

Collection of Adult Health outcomes in GS:

The preclinical questionnaires administered to GS participants before their clinic visit included the Pre-Clinical Questionnaire (PCQ). This was a general health questionnaire asking about physical conditions and symptoms, and lifestyle factors such as diet and smoking. One item on the PCQ, the “Family Health” item asked about the incidence of a variety of serious illnesses in the proband’s family. This item was used to derive all self-reported disease measures, according to the presence/absence of an ‘X’ in the ‘You‘ column of this item.

See <https://www.ed.ac.uk/generation-scotland/our-resources/scottish-family-health-study> for a complete version of the PCQ.

Diagnosis of Major Depressive Disorder was ascertained by a trained researcher using the Structured Clinical Interview for DSM IV (SCID) interview. This took place over the telephone, for participants who had screened positive for suspected depression on the MDQ Mood Questionnaire.

Collection of Physical and Biochemical Measures in GS [1]:

Measurements and samples were taken at a clinic visit by probands. The physical measurements included in the study were:

-	Height (cm), weight (kg)

-	Blood pressure x 2 (Omron BP Monitor), resting pulse

The biochemical measures used were:

-	Total cholesterol (mmol/L)

-	HDL cholesterol (mmol/L)

**Supplementary File 2:** Details of DNA methylation data acquisition and quality control.

Data came from the family-based Generation Scotland: Scottish Family Health Study (GS). GS participants were recruited from GP practices in five regions across Scotland between the years 2006 and 2011 [1]. The probands were aged between 35 and 65 years and were asked to invite first degree relatives to join the study, which had a final sample size of 23, 690. A variety of cognitive, physical, and health data were collected at the study baseline along with blood samples for DNA genotyping.

Blood-based DNA methylation data were obtained on a subset of 5, 200 participants using the Illumina EPIC array. Quality control details have been reported previously Briefly, probes were removed based on: (i) outliers from visual inspection of the log median intensity of the methylated versus unmethylated signal per array; (ii) a beadcount <3 in more than 5% samples and; (iii) ≥5% of samples having a detection p-value >0·05; (iii) any non-autosomal or non-CpG sites, cross-hybridising probes or sites with a SNP at the target CpG or site of single base extension [2]. Samples were removed (i) if there was a mismatch between their predicted sex and recorded sex and/or (ii) if ≥1% of CpGs had a detection p-value >0·05 [3]. We also excluded 3 individuals who answered ‘yes’ to all self-reported disease questions.

As reported in Bermingham et al [3]: “The M-values for CpGs on autosomal chromosomes were pre-corrected for relatedness, estimated blood cell types and processing batch using DISSECT. This was achieved by saving the residuals from a mixed linear model that included methylation as the dependent variable and the following predictor variables: a genetic relatedness matrix fitted in a leave-one-chromosome-out fashion (i.e. SNPs on the same chromosome as the CpG were excluded); proportions of granulocytes, natural killer cells, B-cells, CD4+ T-cells and CD8+ T-cells estimated using the estimateCellCounts function in minfi; and a variable that indicated the batch in which array hybridisation, staining and scanning took place.”

Multi-dimensional scaling was used [4] to generate epigenetic prinicipal co-ordinates on the methylation data after regressing out age, sex, and processing batch. The first 20 PCs were included as covariates to account for additional heterogeneity [3].

For the present analyses, we considered individuals from the DNA methylation subset of GS with birthweight information available. The analysed dataset comprised 841, 753 probes and 1, 395 samples.

For the replication dataset, Illumina HumanMethylationEPIC BeadChips were used to profile genome-wide DNA methylation in whole blood samples from 4, 683 unrelated (<0·05) GS: SFHS participants. These participants were also unrelated (<0·05) to those in the discovery dataset. Quality control of the raw intensity data was carried out using meffil [5] and shinyMethyl [6]. The .idat files were read into R [7] using either the meffil.qc function for the meffil pipeline or minfi’s read.metharray function for shinyMethyl. Quality control was initially carried out using meffil, which was used to remove samples (i) for which a discrepancy between self-reported and methylation-predicted sex (based on the difference between the median copy number intensity for the Y chromosome and the median copy number intensity for the X chromosome) was identified; (ii) that had > 1% CpGs with a detection *p*-value > 0·05; (iii) that showed evidence of dye bias; (iv) that were outliers for the bisulphite conversion control probes; or (v) that had a median methylated signal intensity more than three standard deviations lower than expected. Following the removal of the poor-performing samples detected by meffil, shinyMethyl was used to perform a second round of quality control, as described for the wave 1 dataset. Following the exclusion of samples identified by the steps above, MDS plots were inspected for additional sample outliers and these were excluded too. Poor-performing probes were then identified and removed using meffil. Probes were deemed to have failed if (i) they had a beadcount of < 3 in > 5% samples or (ii) more than > 5% samples had a detection *p*-value > 0·05.

**Supplementary Table 1:** Outputs of minimally adjusted logistic and linear regression models of health traits ~ birth weight residuals + age + sex, with FDR-correction for multiple testing.

| **Trait** | **Odds Ratio** | **95% CI** | **P value** | **P (FDR)** |
| --- | --- | --- | --- | --- |
| SCID depression | 0·854 | 0·78 – 0·94 | 8·5x10^-4^ | 0·0031 |
| SR depression | 0·848 | 0·75 – 0·96 | 0·0067 | 0·018 |
| SR hypertension | 0·885 | 0·75 – 1·04 | 0·14 | 0·19 |
| SR diabetes | 0·921 | 0·72 – 1·18 | 0·521 | 0·52 |
| SR osteoarthritis | 0·762 | 0·61 – 0·96 | 0·019 | 0·041 |
| SR asthma | 0·942 | 0·87 – 1·02 | 0·14 | 0·19 |
| **Trait** | **Beta** | **SE** | **P value** | **P (FDR)** |
| Body Mass Index (kg/m^2^) | 0·068 | 0·015 | 3·9x10^-6^ | 4·3x10^-5^ |
| HDL cholesterol | 0·028 | 0·015 | 0·065 | 0·12 |
| Average systolic BP | -0·013 | 0·013 | 0·31 | 0·34 |
| Average diastolic BP | -0·017 | 0·014 | 0·24 | 0·29 |
| General intelligence (g) | 0·064 | 0·015 | 2·2x10^-5^ | 1·2x10^-4^ |

**Supplementary Table 2:** The 19 CpG sites with P< 1x10^-5^ for their association with birth weight.

| **Probe ID** | **Gene** | **HG19.coordinates** | **Beta** | **P Value** |
| --- | --- | --- | --- | --- |
| cg00966482 | *HERV-FRD* | chr6:11111926 | 0·020 | 6·05x10^-8^ |
| cg06708560 | *DNAJC7/NKIRAS2* | chr17:40170084 | 0·023 | 6·11x10^-7^ |
| cg07593264 | | chr22:43410455 | -0·018 | 8·75x10^-7^ |
| cg27170141 | *CASZ1* | chr1:10711044 | 0·032 | 9·22x10^-7^ |
| cg21931100 | | chr4:116633269 | -0·023 | 1·44x10^-6^ |
| cg00826892 | | chr4:178366013 | 0·032 | 2·75x10^-6^ |
| cg16908601 | *OR2B3* | chr6:29055885 | -0·031 | 2·86x10^-6^ |
| cg15090897 | | chr8:130286226 | -0·022 | 2·91x10^-6^ |
| cg03369398 | | chr12:78973332 | -0·026 | 2·99x10^-6^ |
| cg16639595 | *SRI* | chr7:87856984 | 0·015 | 3·39x10^-6^ |
| cg16300531 | *KSR2* | chr12:118405988 | 0·040 | 5·65x10^-6^ |
| cg19370715 | *CASZ1* | chr1:10710847 | 0·026 | 6·15x10^-6^ |
| cg04988918 | | chr4:178366394 | 0·013 | 6·22x10^-6^ |
| cg08490155 | *SSTR1* | chr14:38676796 | -0·017 | 6·27x10^-6^ |
| cg00590817 | | chr1:8272081 | 0·026 | 8·62x10^-6^ |
| cg01824138 | *CASZ1* | chr1:10699604 | 0·034 | 8·76x10^-6^ |
| cg13524161 | | chr2:200715930 | -0·022 | 8·86x10^-6^ |
| cg16004427 | | chr1:16083101 | 0·028 | 9·19x10^-6^ |
| cg23154832 | *CSMD1* | chr8:4583355 | -0·028 | 9·19x10^-6^ |

**Supplementary Table 3:** CpGs attaining p<1x10^‑5^ in the Replication sample EWAS.

| **Probe ID** | **Gene** | **HG19.coordinates** | **Beta** | **P Value** |
| --- | --- | --- | --- | --- |
| cg04308185 | *ORMDL3* | chr17:38084377 | 0·0811 | 5·4x10^-7^ |
| cg11354629 | *GSX1* | chr13:28366598 | -0·0619 | 1·2x10^-6^ |
| cg19641384 |  | chr2:201694128 | -0·0672 | 2·7x10^-6^ |
| cg19567891 | *LOC254559* | chr15:89921083 | 0·0768 | 2·9x10^-6^ |
| cg22078805 | *FAM171A2* | chr17:42432046 | 0·112 | 3·0x10^-6^ |
| cg02247838 | *CCHCR1* | chr6:31110639 | 0·0402 | 4·0x10^-6^ |
| cg00780250 | *PDS5A* | chr4:39979372 | 0·101 | 4·7x10^-6^ |
| cg24066259 | *CCKBR* | chr11:6281383 | 0·128 | 5·5x10^-6^ |
| cg01503807 | *PARVB* | chr22:44560047 | -0·0525 | 7·0x10^-6^ |
| cg13538398 |  | chr4:129718449 | 0·0384 | 8·0x10^-6^ |
| cg00792008 | *TMEM120B* | chr12:122189621 | 0·0409 | 8·6x10^-6^ |
| cg15942003 | *ORMDL3* | chr17:38084581 | 0·0546 | 9·7x10^-6^ |
| cg23766254 | *FAM171A2* | chr17:42431859 | 0·111 | 9·98x10^-6^ |

**Supplementary Table 4:** CpGs attaining p<1x10^‑5^ in the meta-analysis of discovery sample EWAS and replication sample EWAS. CpG sites and Gene names in bold denote sites that appeared in the discovery sample EWAS.

| **Probe ID** | **Gene** | **HG19.coordinates** | **Effect Size** | **P Value** | **Direction of DNAm across samples** |
| --- | --- | --- | --- | --- | --- |
| **cg00966482** | ***HERV-FRD*** | chr6:11111926 | 0·021 | 5·97x10^-9^ | ++ |
| **cg00590817** |  | chr1:8272081 | 0·028 | 2·84x10^-7^ | ++ |
| cg16365064 |  | chr5:172984486 | 0·021 | 3·24x10^-7^ | ++ |
| **cg27170141** | ***CASZ1*** | chr1:10711044 | 0·031 | 4·37x10^-7^ | ++ |
| cg18321598 |  | chr12:30684738 | 0·017 | 1·03x10^-6^ | ++ |
| **cg00826892** |  | chr4:178366013 | 0·031 | 1·34x10^-6^ | ++ |
| cg26465402 | *ADCY2* | chr5:7803274 | 0·011 | 1·37x10^-6^ | ++ |
| **cg19370715** | ***CASZ1*** | chr1:10710847 | 0·026 | 1·78x10^-6^ | ++ |
| cg02401554 |  | chr4:57203385 | 0·013 | 1·87x10^-6^ | ++ |
| cg08698721 | *MEG3* | chr14:101294147 | -0·015 | 2·24x10^-6^ | -- |
| **cg16639595** | ***SRI*** | chr7:87856984 | 0·015 | 2·34x10^-6^ | ++ |
| cg22178513 | *A1CF* | chr10:52583805 | -0·034 | 2·86x10^-6^ | -- |
| **cg16300531** | ***KSR2*** | chr12:118405988 | 0·04 | 3·02x10^-6^ | ++ |
| **cg03369398** |  | chr12:78973332 | -0·024 | 3·38x10^-6^ | -- |
| cg17293641 | *TNFRSF11B* | chr8:119964144 | 0·02 | 3·41x10^-6^ | ++ |
| cg03425600 |  | chr12:132648585 | 0·014 | 3·76x10^-6^ | ++ |
| cg22572362 | *SLC9A3* | chr5:501938 | 0·026 | 3·84x10^-6^ | ++ |
| cg14809891 |  | chr5:5739504 | -0·021 | 4·10x10^-6^ | -- |
| **cg08490155** | ***SSTR1*** | chr14:38676796 | -0·017 | 4·56x10^-6^ | -- |
| **cg01824138** | ***CASZ1*** | chr1:10699604 | 0·032 | 4·65x10^-6^ | ++ |
| cg14534277 | *DSC3* | chr18:28623874 | -0·022 | 5·04x10^-6^ | -- |
| cg00296348 |  | chr10:134721479 | 0·024 | 5·13x10^-6^ | ++ |
| cg19706390 | *CASZ1* | chr1:10709702 | 0·036 | 5·47x10^-6^ | ++ |
| cg18907109 | *HEPACAM2* | chr7:92849682 | -0·01 | 5·57x10^-6^ | -- |
| **cg04988918** |  | chr4:178366394 | 0·013 | 5·72x10^-6^ | +? |
| **cg21931100** |  | chr4:116633269 | -0·02 | 6·35x10^-6^ | -- |
| cg24248329 | *NFYC* | chr1:41175132 | 0·022 | 6·82x10^-6^ | ++ |
| cg03835140 | *NKPD1* | chr19:45662154 | 0·015 | 7·08x10^-6^ | ++ |
| cg03964940 | *STX11* | chr6:144477779 | 0·022 | 7·24x10^-6^ | ++ |
| cg01152056 |  | chr6:166260319 | 0·038 | 7·54x10^-6^ | ++ |
| **cg07593264** |  | chr22:43410455 | -0·016 | 7·60x10^-6^ | -+ |
| cg09363850 |  | chr11:58090434 | -0·029 | 8·03x10^-6^ | -- |
| cg03721657 | *HCCA2* | chr11:1571033 | 0·016 | 8·65x10^-6^ | ++ |
| cg20014974 |  | chr1:8271918 | 0·014 | 8·75x10^-6^ | ++ |
| cg26086468 |  | chr18:5628160 | -0·014 | 9·90x10^-6^ | -- |
| cg19863411 | *PCYT2* | chr17:79869801 | 0·022 | 9·98x10^-6^ | ++ |

**Supplementary Table 5:** Sensitivity analysis run on CpGs attaining P<1x10^-5^ in the discovery sample, with % change in effect size (Beta) from original analysis.

| **Probe ID** | **Gene** | **HG19.coordinates** | **Beta** | **P Value** | **Beta % change** |
| --- | --- | --- | --- | --- | --- |
| cg00966482 | *HERV-FRD* | chr6:11111926 | 0·019 | 1·20x10^-6^ | 5·04 |
| cg06708560 | *DNAJC7/NKIRAS2* | chr17:40170084 | 0·023 | 2·29x10^-6^ | -1·16 |
| cg07593264 |  | chr22:43410455 | -0·016 | 3·65x10^-5^ | 11·12 |
| cg27170141 | *CASZ1* | chr1:10711044 | 0·032 | 6·70x10^-6^ | 1·23 |
| cg21931100 |  | chr4:116633269 | -0·023 | 4·99x10^-6^ | -0·95 |
| cg00826892 |  | chr4:178366013 | 0·034 | 2·06x10^-6^ | -6·79 |
| cg16908601 | *OR2B3* | chr6:29055885 | -0·030 | 1·21x10^-5^ | 1·07 |
| cg15090897 |  | chr8:130286226 | -0·021 | 2·49x10^-5^ | 6·26 |
| cg03369398 |  | chr12:78973332 | -0·025 | 2·82x10^-5^ | 4·61 |
| cg16639595 | *SRI* | chr7:87856984 | 0·014 | 2·49x10^-5^ | 3·84 |
| cg16300531 | *KSR2* | chr12:118405988 | 0·041 | 1·64x10^-5^ | -0·77 |
| cg19370715 | *CASZ1* | chr1:10710847 | 0·024 | 8·91x10^-5^ | 7·25 |
| cg04988918 |  | chr4:178366394 | 0·014 | 3·64x10^-6^ | -8·51 |
| cg08490155 | *SSTR1* | chr14:38676796 | -0·017 | 4·26x10^-5^ | 2·46 |
| cg00590817 |  | chr1:8272081 | 0·029 | 2·00x10^-6^ | -12·67 |
| cg01824138 | *CASZ1* | chr1:10699604 | 0·034 | 2·78x10^-5^ | -0·68 |
| cg13524161 |  | chr2:200715930 | -0·021 | 7·30x10^-5^ | 5·53 |
| cg16004427 |  | chr1:16083101 | 0·025 | 1·00x10^-4^ | 7·40 |
| cg23154832 | *CSMD1* | chr8:4583355 | -0·028 | 3·23x10^-5^ | 1·40 |

**Supplementary Table 6:** Significant (P<5x10-8) GWAS catalogue outputs for SNPs in genes that mapped to the CpG sites from the EWAS with P<1x10-5

| **Gene** | **Trait** | **SNP** | **P-Value** | **Study Link** |
| --- | --- | --- | --- | --- |
| *CASZ1* | Systolic blood pressure | rs880315 | 7·00E-18 | www.ncbi.nlm.nih.gov/pubmed/29403010 |
| *CASZ1* | Diastolic blood pressure | rs880315 | 6·00E-12 | www.ncbi.nlm.nih.gov/pubmed/29403010 |
| *CASZ1* | Blood pressure | rs880315 | 3·00E-10 | www.ncbi.nlm.nih.gov/pubmed/21572416 |
| *CASZ1* | Systolic blood pressure | rs880315 | 6·00E-10 | www.ncbi.nlm.nih.gov/pubmed/25249183 |
| *CASZ1* | Hypertension | rs880315 | 2·00E-09 | www.ncbi.nlm.nih.gov/pubmed/25249183 |
| *CASZ1* | Diastolic blood pressure | rs880315 | 4·00E-10 | www.ncbi.nlm.nih.gov/pubmed/28739976 |
| *CASZ1* | Systolic blood pressure | rs880315 | 9·00E-16 | www.ncbi.nlm.nih.gov/pubmed/28739976 |
| *CASZ1* | Pulse pressure | rs880315 | 2·00E-10 | www.ncbi.nlm.nih.gov/pubmed/28739976 |
| *CASZ1* | Mean arterial pressure | rs880315 | 5·00E-17 | www.ncbi.nlm.nih.gov/pubmed/27618448 |
| *CASZ1* | Systolic blood pressure | rs880315 | 2·00E-14 | www.ncbi.nlm.nih.gov/pubmed/27618452 |
| *CASZ1* | Diastolic blood pressure | rs880315 | 1·00E-11 | www.ncbi.nlm.nih.gov/pubmed/27618452 |
| *CASZ1* | Pulse pressure | rs880315 | 5·00E-09 | www.ncbi.nlm.nih.gov/pubmed/29403010 |
| *CASZ1* | Mean arterial pressure | rs880315 | 2·00E-16 | www.ncbi.nlm.nih.gov/pubmed/29403010 |
| *CASZ1* | Atrial fibrillation | rs880315 | 5·00E-09 | www.ncbi.nlm.nih.gov/pubmed/29892015 |
| *CASZ1* | Systolic blood pressure x alcohol consumption interaction (2df test) | rs17035646 | 1·00E-36 | www.ncbi.nlm.nih.gov/pubmed/29912962 |
| *CASZ1* | Systolic blood pressure x alcohol consumption interaction (2df test) | rs35295665 | 5·00E-27 | www.ncbi.nlm.nih.gov/pubmed/29912962 |
| *CASZ1* | Systolic blood pressure x alcohol consumption (light vs heavy) interaction (2df test) | rs17035646 | 7·00E-14 | www.ncbi.nlm.nih.gov/pubmed/29912962 |
| *CASZ1* | Systolic blood pressure x alcohol consumption (light vs heavy) interaction (2df test) | rs17035646 | 9·00E-13 | www.ncbi.nlm.nih.gov/pubmed/29912962 |
| *CASZ1* | Diastolic blood pressure x alcohol consumption interaction (2df test) | rs34071855 | 5·00E-29 | www.ncbi.nlm.nih.gov/pubmed/29912962 |
| *CASZ1* | Diastolic blood pressure x alcohol consumption interaction (2df test) | rs34071855 | 2·00E-26 | www.ncbi.nlm.nih.gov/pubmed/29912962 |
| *CASZ1* | Diastolic blood pressure | rs880315 | 6·00E-16 | www.ncbi.nlm.nih.gov/pubmed/27618447 |
| *CASZ1* | Stroke | rs880315 | 4·00E-10 | www.ncbi.nlm.nih.gov/pubmed/29531354 |
| *CASZ1* | Ischemic stroke | rs880315 | 6·00E-09 | www.ncbi.nlm.nih.gov/pubmed/29531354 |
| *CASZ1* | Diastolic blood pressure x alcohol consumption (light vs heavy) interaction (2df test) | rs17035646 | 1·00E-11 | www.ncbi.nlm.nih.gov/pubmed/29912962 |
| *CASZ1* | Pulse pressure x alcohol consumption interaction (2df test) | rs35295665 | 9·00E-17 | www.ncbi.nlm.nih.gov/pubmed/29912962 |
| *CASZ1* | Pulse pressure x alcohol consumption interaction (2df test) | rs17035646 | 2·00E-20 | www.ncbi.nlm.nih.gov/pubmed/29912962 |
| *CASZ1* | Mean arterial pressure x alcohol consumption interaction (2df test) | rs34071855 | 4·00E-20 | www.ncbi.nlm.nih.gov/pubmed/29912962 |
| *CASZ1* | Mean arterial pressure x alcohol consumption (light vs heavy) interaction (2df test) | rs17035646 | 5·00E-14 | www.ncbi.nlm.nih.gov/pubmed/29912962 |
| *CASZ1* | Atrial fibrillation | rs284277 | 1·00E-09 | www.ncbi.nlm.nih.gov/pubmed/30061737 |
| *CASZ1* | Diastolic blood pressure (cigarette smoking interaction) | rs880315 | 7·00E-42 | www.ncbi.nlm.nih.gov/pubmed/29455858 |
| *CASZ1* | Systolic blood pressure (cigarette smoking interaction) | rs880315 | 2·00E-54 | www.ncbi.nlm.nih.gov/pubmed/29455858 |
| *CASZ1* | Male-pattern baldness | rs2242288 | 3·00E-10 | www.ncbi.nlm.nih.gov/pubmed/28196072 |
| *CASZ1* | Male-pattern baldness | rs59304342 | 4·00E-08 | www.ncbi.nlm.nih.gov/pubmed/28196072 |
| *CSMD1* | Schizophrenia | rs10503253 | 2·00E-08 | www.ncbi.nlm.nih.gov/pubmed/21926974 |
| *CSMD1* | Cannabis dependence symptom count | rs77378271 | 2·00E-08 | www.ncbi.nlm.nih.gov/pubmed/27028160 |
| *CSMD1* | Autism spectrum disorder, attention deficit-hyperactivity disorder, bipolar disorder, major depressive disorder, and schizophrenia (combined) | rs10503253 | 4·00E-08 | www.ncbi.nlm.nih.gov/pubmed/23453885 |
| *CSMD1* | Schizophrenia | rs10503253 | 1·00E-08 | www.ncbi.nlm.nih.gov/pubmed/25056061 |
| *CSMD1* | Menarche (age at onset) | rs2688325 | 2·00E-09 | www.ncbi.nlm.nih.gov/pubmed/25231870 |
| *CSMD1* | Menarche (age at onset) | rs7828501 | 1·00E-13 | www.ncbi.nlm.nih.gov/pubmed/25231870 |
| *CSMD1* | Menarche (age at onset) | rs7463166 | 1·00E-08 | www.ncbi.nlm.nih.gov/pubmed/25231870 |
| *CSMD1* | Resting heart rate | rs145669495 | 2·00E-08 | www.ncbi.nlm.nih.gov/pubmed/28270201 |
| *CSMD1* | Hand grip strength | rs752045 | 5·00E-10 | www.ncbi.nlm.nih.gov/pubmed/27325353 |
| *CSMD1* | Schizophrenia | rs13261217 | 2·00E-09 | www.ncbi.nlm.nih.gov/pubmed/28991256 |
| *CSMD1* | Urinary albumin excretion | rs55798132 | 3·00E-08 | www.ncbi.nlm.nih.gov/pubmed/30220432 |
| *CSMD1* | Schizophrenia | rs139425113 | 8·00E-09 | www.ncbi.nlm.nih.gov/pubmed/29483656 |
| *KSR2* | Coronary artery disease | rs11830157 | 2·00E-09 | www.ncbi.nlm.nih.gov/pubmed/26343387 |
| *KSR2* | Depression | rs7973260 | 2·00E-08 | www.ncbi.nlm.nih.gov/pubmed/27089181 |
| *KSR2* | Post bronchodilator FEV1/FVC ratio | rs183032784 | 6·00E-09 | www.ncbi.nlm.nih.gov/pubmed/26634245 |
| *OR2B3* | Autism spectrum disorder or schizophrenia | rs144649399 | 1·00E-12 | www.ncbi.nlm.nih.gov/pubmed/28540026 |
| *OR2B3* | Autism spectrum disorder or schizophrenia | rs115661163 | 1·00E-11 | www.ncbi.nlm.nih.gov/pubmed/28540026 |
| *OR2B3* | Autism spectrum disorder or schizophrenia | rs115329265 | 2·00E-27 | www.ncbi.nlm.nih.gov/pubmed/28540026 |
| *OR2B3* | Autism spectrum disorder or schizophrenia | rs116663187 | 2·00E-15 | www.ncbi.nlm.nih.gov/pubmed/28540026 |
| *OR2B3* | Autism spectrum disorder or schizophrenia | rs116137698 | 3·00E-26 | www.ncbi.nlm.nih.gov/pubmed/28540026 |
| *OR2B3* | Autism spectrum disorder or schizophrenia | rs144911693 | 5·00E-14 | www.ncbi.nlm.nih.gov/pubmed/28540026 |
| *OR2B3* | Autism spectrum disorder or schizophrenia | rs1150688 | 8·00E-11 | www.ncbi.nlm.nih.gov/pubmed/28540026 |
| *OR2B3* | Autism spectrum disorder or schizophrenia | rs151267808 | 1·00E-22 | www.ncbi.nlm.nih.gov/pubmed/28540026 |
| *OR2B3* | Autism spectrum disorder or schizophrenia | rs115123779 | 1·00E-17 | www.ncbi.nlm.nih.gov/pubmed/28540026 |
| *OR2B3* | Blood protein levels | rs3129682 | 1·00E-19 | www.ncbi.nlm.nih.gov/pubmed/28240269 |
| *OR2B3* | Blood protein levels | rs3131085 | 1·00E-09 | www.ncbi.nlm.nih.gov/pubmed/28240269 |
